# Inhibition of BET family proteins suppresses African swine fever virus infection

**DOI:** 10.1101/2021.11.02.467039

**Authors:** Yaru Zhao, Qingli Niu, Saixia Yang, Jifei Yang, Zhonghui Zhang, Shuxian Geng, Jie Fan, Zhijie Liu, Guiquan Guan, Zhiqing Liu, Jia Zhou, Haitao Hu, Jianxun Luo, Hong Yin

## Abstract

African swine fever (ASF), an acute, severe, highly contagious disease caused by African swine fever virus (ASFV) infection in domestic pigs and boars, has a mortality rate of up to 100%. Because effective vaccines and treatments for ASF are lacking, effective control of the spread of ASF remains a great challenge for the pig industry. Host epigenetic regulation is essential for the viral gene transcription. Bromodomain and extraterminal (BET) family proteins, including BRD2, BRD3, BRD4, and BRDT, are epigenetic “readers” critical for gene transcription regulation. Among these proteins, BRD4 recognizes acetylated histones via its two bromodomains (BD1 and BD2) and recruits transcription factors, thereby playing a pivotal role in transcriptional regulation and chromatin remodeling during viral infection. However, how BET/BRD4 regulates ASFV replication and gene transcription is unknown. Here, we randomly selected 12 representative BET family inhibitors and compared their effects on ASFV infection in pig’s primary alveolar macrophages (PAMs). They were found to inhibit viral infection by interfering with the different stages of viral life cycle (attachment, internalization, desencapsidation and formation of viral factories). The four most effective inhibitors (ARV-825, ZL0580, I-BET-762 and PLX51107) were selected for further antiviral activity analysis. These BET/BRD4 inhibitors dose-dependently decreased the ASFV titer, viral RNA transcription and protein production in PAMs. Collectively,our study reported novel activity of BET/BRD4 inhibitors in inducing suppression of ASFV infection, providing insights into role of BET/BRD4 in epigenetic regulation of ASFV and potential new strategies for ASF prevention and control.

**IMPORTANCE:** Since the continuing spread of the ASFV in the world, and lack of commercial vaccines, the development of improved control strategies including antiviral drugs are urgently needed. BRD4 is an important epigenetic factor and has been commonly used for drug development for tumor treatment. Furthermore, the latest research showed that BET/BRD4 inhibition could suppress replication of virus. In this study, we first showed the inhibitory effect of agents targeting BET/BRD4 on ASFV infection with no significant host cytotoxicity. Then, we found 4 BET/BRD4 inhibitors which can inhibit ASFV replication, RNA transcription and protein synthesis. Finally, we analyzed 4 inhibitors’ biological effect on BRD4 according to the structure of BRD4, and docking analysis of BET-762, PLX51107, ARV-825 and ZL0580 binding to BD1 and BD2 domains of BRD4 was performed. Our findings support the hypothesis that BET/BRD4 can be considered as attractive host targets in antiviral drug discovery against ASFV.

## INTRODUCTION

African swine fever (ASF), a highly contagious viral disease in swine infected with African swine fever virus (ASFV), exhibits a high mortality rate approaching 100% and has severe economic consequences for affected countries (1). ASF is clinically characterized by high fever, spotty skin, cyanosis, extensive bleeding of the internal organs, and disturbance of the respiratory and nervous systems (2). ASF was first introduced to Liaoning Province of China in August 2018, when genotype II ASFV resulted in numerous outbreaks within domestic pigs (http://www.oie.int/). These outbreaks resulted in economic losses of several billion dollars to China’s pig breeding industry and national economy, seriously affecting the lives of Chinese residents, national economic development and the pig industry (3).

The ASFV genome is large and complex, and the mechanism by which replication is regulated is unclear thus far. Although ASFV was discovered nearly a hundred years ago, no commercial vaccines or cost-effective antiviral drugs are available to effectively prevent ASF worldwide. ASFV, a tick-borne, double-stranded DNA virus and the only member of the *Asfarviridae* family, genus *Asfivirus*, mainly infects myeloid lineage cells, especially monocytes/macrophages and dendritic cells (4). The replication of ASFV primarily occurs in the cytoplasm, but a transient nucleation progress occurs at the early stage (5, 6). However, the nuclear replication mechanism is not clear. ASFV encodes more than 200 proteins, including at least 54 structural proteins and more than 100 nonstructural proteins, which are involved in replication of the genome and assembly of the virion, respectively, and also regulate host cell function and immune evasion (7). Following viral infection, the host cell transcriptional machinery is required for viral gene expression, indicating involvement of host epigenetic factors in this process (8).

Histone lysine acetylation is a key mechanism in chromatin processes and the regulation of gene transcription (9). BET family proteins include BRD2, BRD3, BRD4, and BRDT, which have important biological functions, such as their ability to mediate transcriptional regulation and chromatin remodeling (10). BRD4 recruits positive transcriptional elongation factor (P-TEFb) complex, which plays an essential role in transcriptional regulation by RNA polymerase II (RNA Pol II) in eukaryotes (11). BRD4 is one of the most important proteins in the BET family, and contains two bromo-domains (BD1 and BD2). BRD4 is not only a chromatin reader protein but also an epigenetic regulatory factor and transcription cofactor closely related to gene transcription, the cell cycle and apoptosis (12, 13). Abnormal BRD4 protein expression can lead to the disordered expression of various genes and thus affects the function of related genes. BRD4 also plays an important role in DNA replication, transcription and repair (14). Among host molecules, BRD4 can be used by DNA viruses to regulate the transcription of viral genes during viral replication through critical protein-protein interactions. BRD4 and its inhibitors have been widely studied as potential antitumor therapies. The latest research showed that BRD4 inhibition activated the cGAS-STING pathway of the antiviral innate immune response by leading to DNA damage-dependent signaling and attenuated viral attachment of pseudorabies virus (PRV) (15). In addition, a BRD4 inhibitor was found to suppress human immunodeficiency virus (HIV) by inhibiting Tat transactivation and transcription elongation and by inducing a repressive chromatin structure at the HIV promoter (16).

However, potential effect of BET/BRD4 on ASFV replication and viral transcription has not been evaluated. ASFV may alter the epigenetic status of host chromatin to modulate cellular gene expression for its own benefit. Therefore, we focused on the biological effects of representative BET/BRD4 inhibitors on the replication and transcription of ASFV *in vitro*, and our results may open new avenues for the effective prevention and control of ASF.

## RESULTS

### Cytotoxicity of BET inhibitors in PAMs

The inhibitors were used at nine different concentrations ranging from 0.5 µM to 240 µM to evaluate cytotoxicity by using the CCK-8 assay. The results indicated that at least 7 of the inhibitors did not cause a significant increase in cell death, and the cell viability reached more than 60% when the concentrations of the inhibitors were up to 20 µM, but significant cytotoxic effects on the PAMs were observed at concentrations from 40 µM to 240 µM. Cell viability was still more than 50% when the inhibitors INCB054329 and CPI-203 were used at 80 µM. The inhibitors demonstrated potent cytotoxic effects on PAMs at 10 μM (ARV-825 and AZD5153), 20 μM (PLX51107, PFI-1, RVX-208, ZL0580 and (+)-JQ1), 40 μM (OTX051, MS436 and I-BET-762) and 80 μM (INCB054329 and CPI-203) (Figure S2). The organic solvent DMSO had no cytotoxic effect on PAMs (data not shown). In summary, even though most of these BET inhibitors are commercially available as research tools, some of them show a certain degree of cytotoxicity against PAMs at high concentrations at which the primary cells are more sensitive to the inhibitors.

### Effect of BET inhibitors on ASFV transcription in PAMs

To determine whether the BET inhibitors could affect ASFV gene transcription by altering the functions of BET proteins, a time-of-addition assay was conducted to evaluate the effects of 12 BET/BRD4 inhibitors on specific step(s) of the ASFV life cycle. Cells were treated with the individual BET inhibitors at 5 µM, and the functional role of BET in BET inhibitor-induced ASFV gene transcription was evaluated using real-time PCR. The relative expression levels of *CP204L* (early), *B646L* (late) and *GAPDH* in the cells treated with individual BET inhibitors were measured and compared with those in the control (DMSO; negative control [NC]) group. Pretreatment with the BET inhibitors potently suppressed ASFV gene transcription in the cells (Figure 1A). A significant inhibitory effect on transcription of the *CP204L* gene, which is expressed early during the ASFV infection cycle, was observed when the inhibitors were applied simultaneously with ASFV infection, but the effect was less pronounced than that observed upon pretreatment (Figure 1B). Moreover, neither *CP204L* nor *B646L* gene transcription was inhibited by BET inhibitor post-treatment (Figure 1C). Interestingly, pretreatment with PLX51107 and ZL0580 almost completely inhibited ASFV gene transcription. Since accumulating evidence suggests that BET/BRD4 plays an important role in regulating viral transcriptional (17-19) and based on our above results, four representative inhibitors (PLX51107, I-BET762, ZL0580 and ARV-825) with the greatest inhibitory effects under both pretreatment and cotreatment conditions were selected for further experiments. Among these four inhibitors, the first two (PLX51107 and I-BET762) are broad-spectrum BET family inhibitors, while the latter two (ZL0580 and ARV-825) are BRD4-specific inhibitors. Collectively, these results suggest that BET/BRD4 inhibition results in decreased ASFV gene transcription in ASFV infection, and the expression of ASFV *CP204L* and *B646L* upon inhibitor treatment significantly differed from that in untreated cells (DMSO-treated group) *in vitro*. Thus, further experiments were performed.

**Figure 1:**
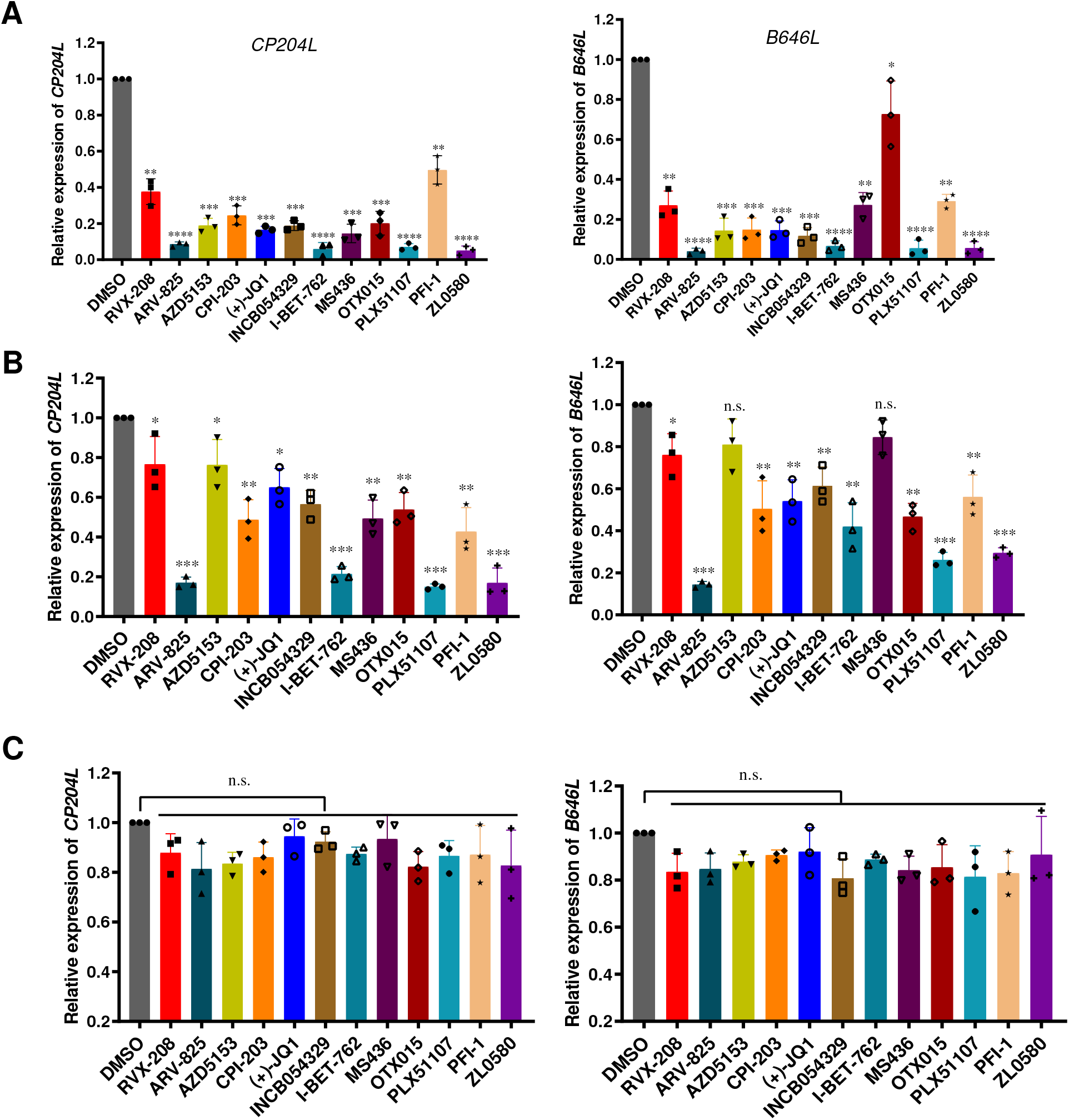
Effect of BET family inhibitors on ASFV gene transcription. (A) Expression levels of *CP204L* and *B646L* mRNA from ASFV after pretreatment, (B) cotreatment and (C) posttreatment with the inhibitors were detected by real-time PCR. Data were normalized to data from the DMSO-treated samples. PAMs in 12-well plates were pretreated, cotreated or posttreated with individual inhibitors or DMSO in relation to ASFV (MOI =0.1) infection. The samples were collected at 24 h postinfection under pre- and cotreatment conditions with the inhibitors. For the posttreatment samples, the cells were first infected with ASFV for 4 h, followed by inhibitor treatment for 16 h, after which the samples were collected. The concentration of the BET/BRD4 inhibitors was 5 μM. Error bars show the SD of replicates qPCR experiments. All experiments were independently conducted at least 3 times. Statistical significance is denoted by \**P* < 0.05, \*\**P* < 0.01, \*\*\**P* < 0.001, \*\*\*\**P* < 0.0001.

### PLX51107, I-BET762, ZL0580 and ARV-825 inhibit viral infection in a time-dependent manner

The structures of ARV-825, ZL0580, I-BET-762 and PLX51107 are shown in Figure 2A. The CC_50_ values, the concentrations of the 4 inhibitors at which they caused 50% cell death, were calculated in PAMs. The CC_50_ values of ARV-825, ZL0580, I-BET-762 and PLX51107 were determined to be 10.11 μM (95% CI = 9.18-11.11), 25.3 μM (95% CI = 21.57-31.11), 35.86 μM (95% CI = 25.38-86.79) and 19.37 μM (95% CI = 15.91-24.68), respectively (Figure 2B). In antiviral experiments, to mitigate their cytotoxic effects, ARV-825, ZL0580, PLX51107 and I-BET-762 were used at maximum concentrations of 1 μM, 10 μM, 5 μM and 10 μM, respectively, which were lower than the CC_50_ values. The duration over which the 4 inhibitors inhibited the replication of ASFV was further evaluated. The four inhibitors were added to PAM culture medium for 16 h prior to ASFV infection. Samples were collected at 4 h, 12 h, 24 h and 48 h after infection. Relative expression of the *B646L* gene was then detected by real-time PCR. The results indicated inhibitory effects to different degrees depending on the inhibitor. The inhibitory effects of ARV-825 and I-BET-762 were observed at 12 h after ASFV infection, while a strong antiviral effect at the early stages of replication that continued for 48 h was observed after ZL0580 treatment (Figure 3). PLX51107 also exerted a time-dependent inhibitory effect. These results indicate that these 4 inhibitors significantly inhibit the replication of ASFV at the early, middle and late stages of ASFV infection, although they inhibit the functions of BET/BRD4 in different manners.

**Figure 2:**
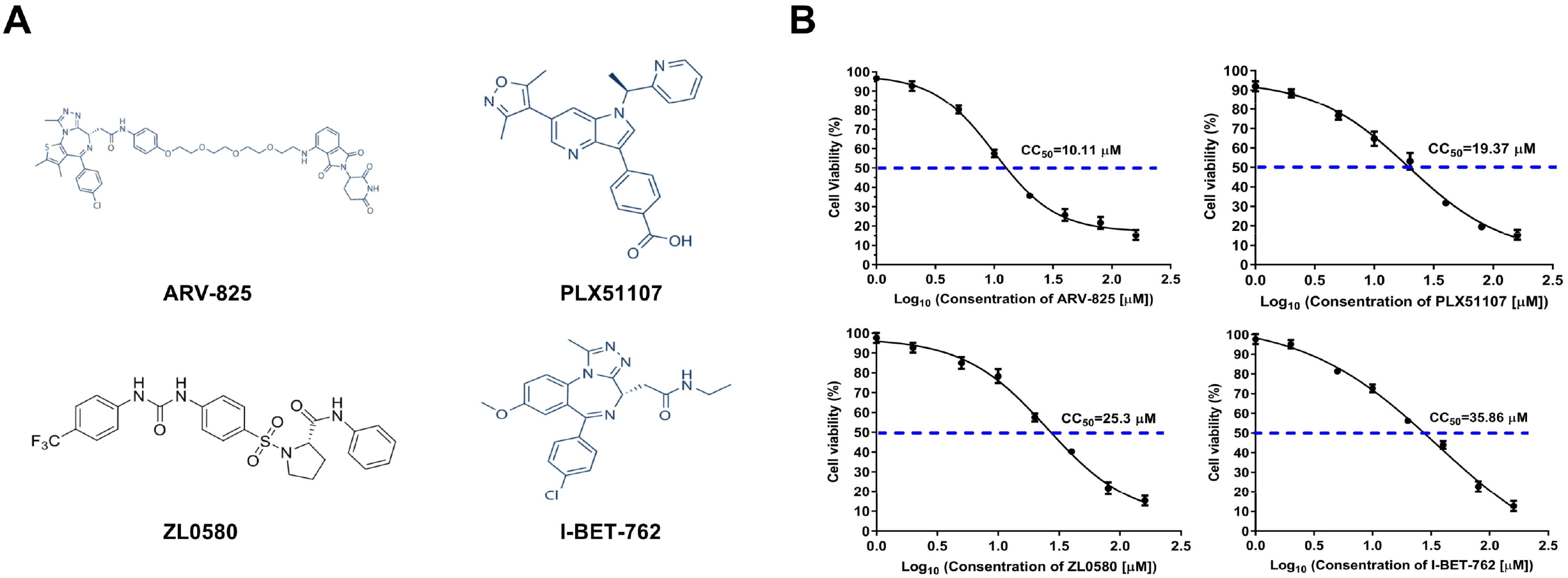
Structures of inhibitors and CC_50_ values. (A) The structures of ARV-825, ZL0580, I-BET-762 and PLX51107. (B) The CC_50_ values of ARV-825, ZL0580, I-BET-762 and PLX51107 against PAMs were calculated.

**Figure 3:**
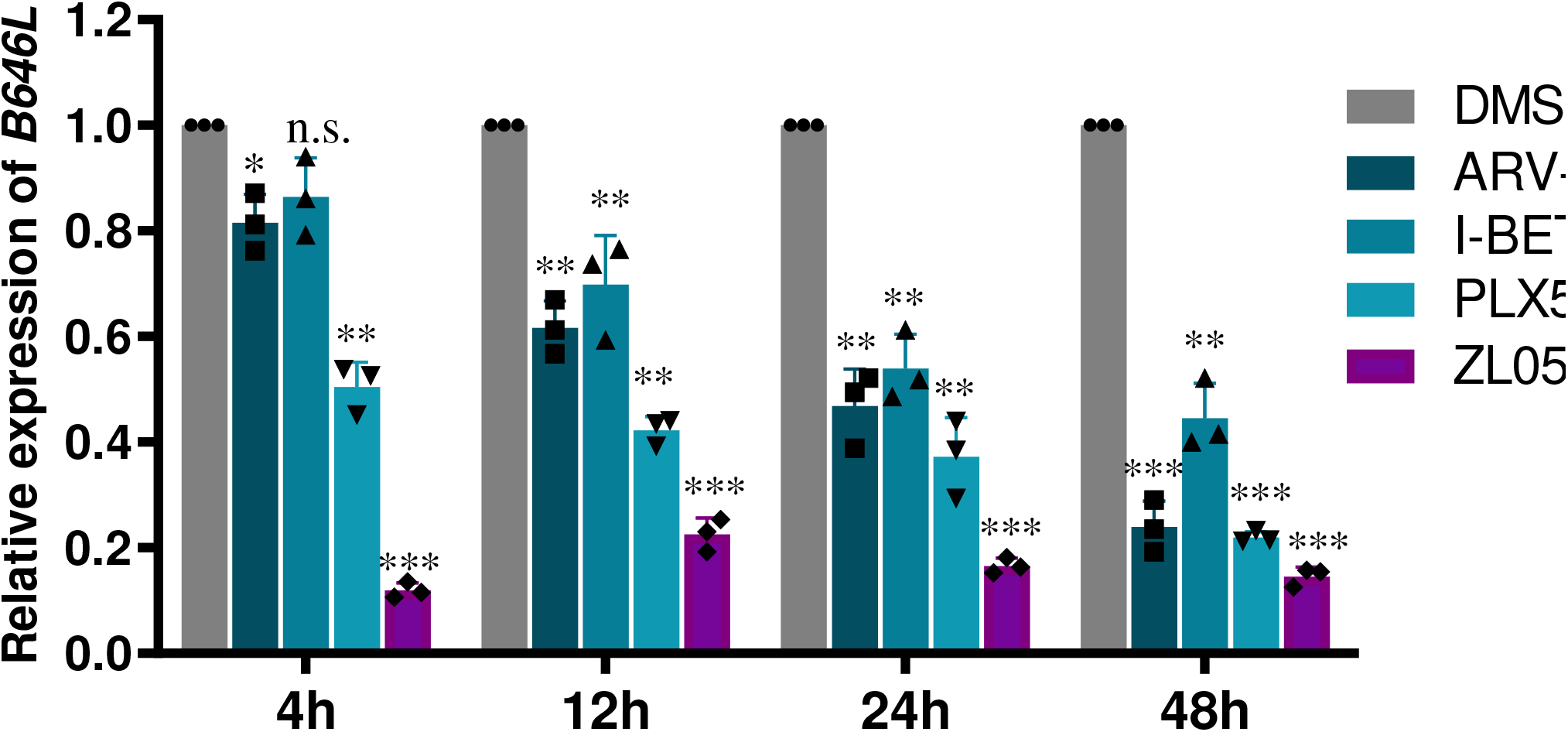
Time-dependent effect of 4 inhibitors on PAMs. The four inhibitors act throughout the whole ASFV infection cycle to decrease ASFV RNA levels. PAMs were treated with 1 μM ARV-825, 10 μM ZL0580, 10 μM I-BET-762 and 5 μM PLX51107 for 16 h prior to ASFV infection (MOI =0.1). ASFV *B646L* mRNA levels at 4 h, 12, 24 h and 48 h postinfection were detected and analyzed by RT-qPCR. Data were normalized to data from DMSO-treated samples. Statistical significance is denoted by \**P* < 0.05, \*\**P* < 0.01, \*\*\**P* < 0.001.

### Inhibitory effect of BET/BRD4 on ASFV infection of PAMs in a dose-dependent manner

In an ASFV suppression model, PAMs were treated with 4 individual inhibitors, and their potential dose-dependent antiviral activity against ASFV was evaluated. We treated ASFV-infected PAMs with the individual inhibitors at increasing concentrations from 0.1 μM to 10 μM depending on the inhibitor. As shown in Figure 4A, the viral yields decreased significantly from 6 to 1.6 log HAD_50_/ml at a concentration of 1 μM (ARV-825), 5 μM (PLX51107) or 10 μM (ZL0580 and I-BET-762) (*P* < 0.05 or 0.001). At the gene transcription level, ZL0580 did not significantly suppress late ASFV *B646L* mRNA expression at concentrations lower than 2 μM. All four inhibitors clearly suppressed the early expression of ASFV *CP204L* mRNA (*P* < 0.05, 0.001 or 0.0001) (Figure 4B). Importantly, further analysis of protein expression levels revealed that viral p72 protein expression levels were clearly suppressed in ASFV-infected PAMs treated with the 4 inhibitors in a dose-dependent manner, especially upon treatment with 0.25-1 μM ARV-825, which fully inhibited expression of the p72 protein (Figure 4C). These results indicated that the four BET/BRD4 inhibitors suppressed the ASFV titer, as well as mRNA and protein synthesis during replication. Based on these results, maximum concentrations of 1 μM (ARV-825), 10 μM (ZL0580 and I-BET-762) and 5 μM (PLX51107) were selected for further evaluation of the effects of the inhibitors against ASFV infection.

**Figure 4:**
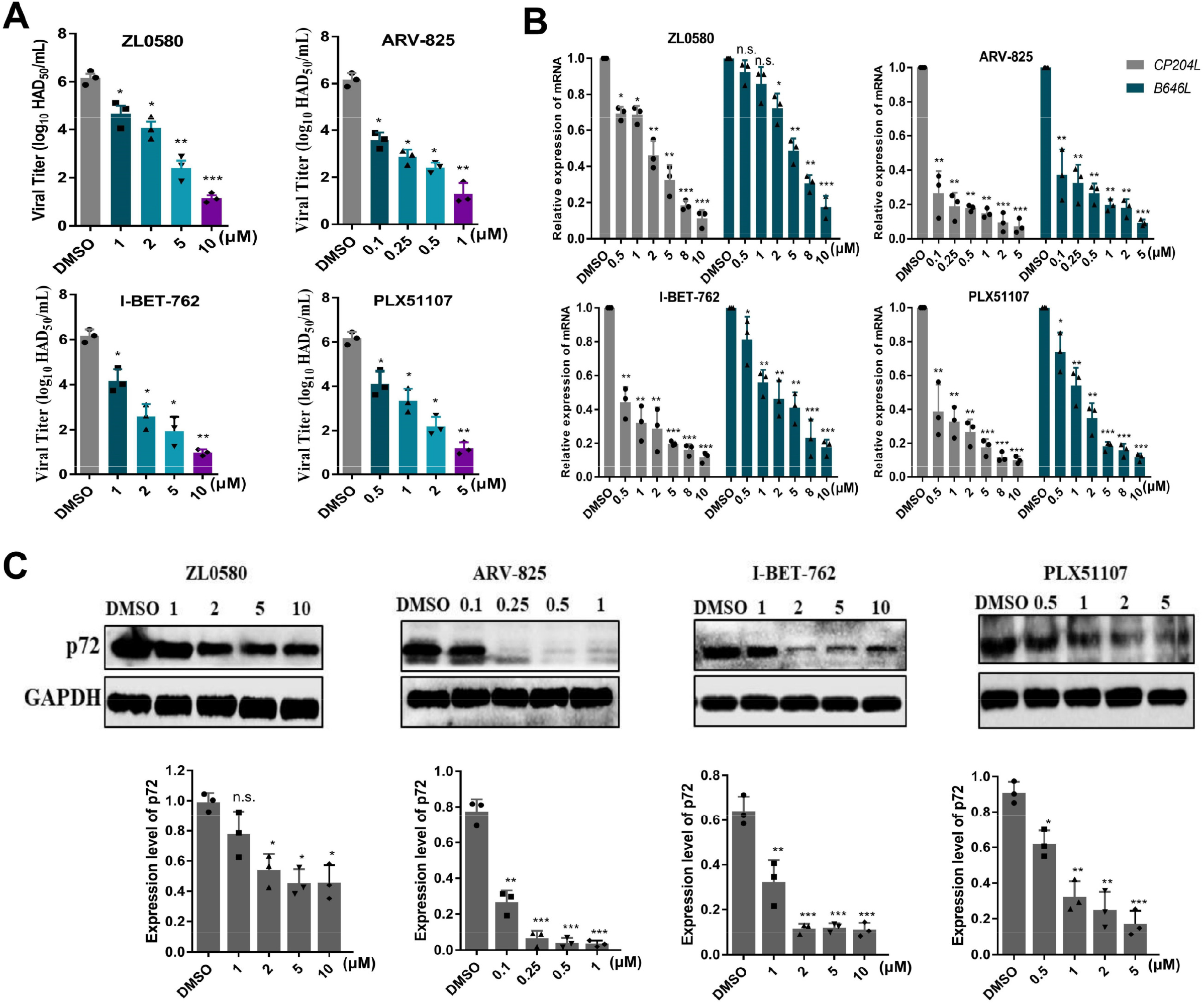
Dose-dependent effects of ARV-825, I-BET-762, PLX51107 and ZL0580 on ASFV replication. (A) ASFV yield in PAMs decreased significantly in a dose-dependent manner with 4 inhibitor pretreatment. (B) ASFV *CP204L* and *B646L* mRNA levels were analyzed by RT qPCR. (C) The expression of p72 in the presence of 4 inhibitors at several concentrations was evaluated by WB analysis. PAMs in 12-well plates were treated with individual inhibitors or DMSO for 16 h prior to ASFV infection (MOI =0.1). The samples were collected at 24 h postinfection. Data were normalized to data from DMSO-treated samples. Error bars show the SD of replicate qPCR experiments. All experiments were independently conducted at least 3 times. Statistical significance is denoted by \**P* < 0.05, \*\**P* < 0.01, \*\*\**P* < 0.001.

### BRD4 Inhibition attenuates viral infection

To determine the influence for viral infection by BRD4 inhibition, we pre-treated PAM cells with DMSO or ARV-825 for 2 h at 37°C. Then, the cells were incubated with R18 labeled ASFV and BRD4 inhibitors for 1 h at 4°C. Immunofluorescence analysis indicated that the fluorescent signals in cells treated with DMSO were stronger than that in cells treated with ARV-825, thus suggesting that BRD4 inhibitors influenced ASFV attachment (Figure 5A). CD2v is outer envelope protein involved in viral attachment. Similar phenomena were also observed in determination of CD2v protein expression by Western blotting, CD2v expressed level was significantly decreased in the cells with inhibitors treatment (Figure 5B). Furthermore, we found that the viral internalization, desencapsidation and factories in infected cells with different treated time points were also affected by BRD4 inhibition. Clear fluorescent signals in cells treated with DMSO were observed and weaker signals were found in cells treated with ARV-825 (Figure 5C-E).

**Figure 5:**
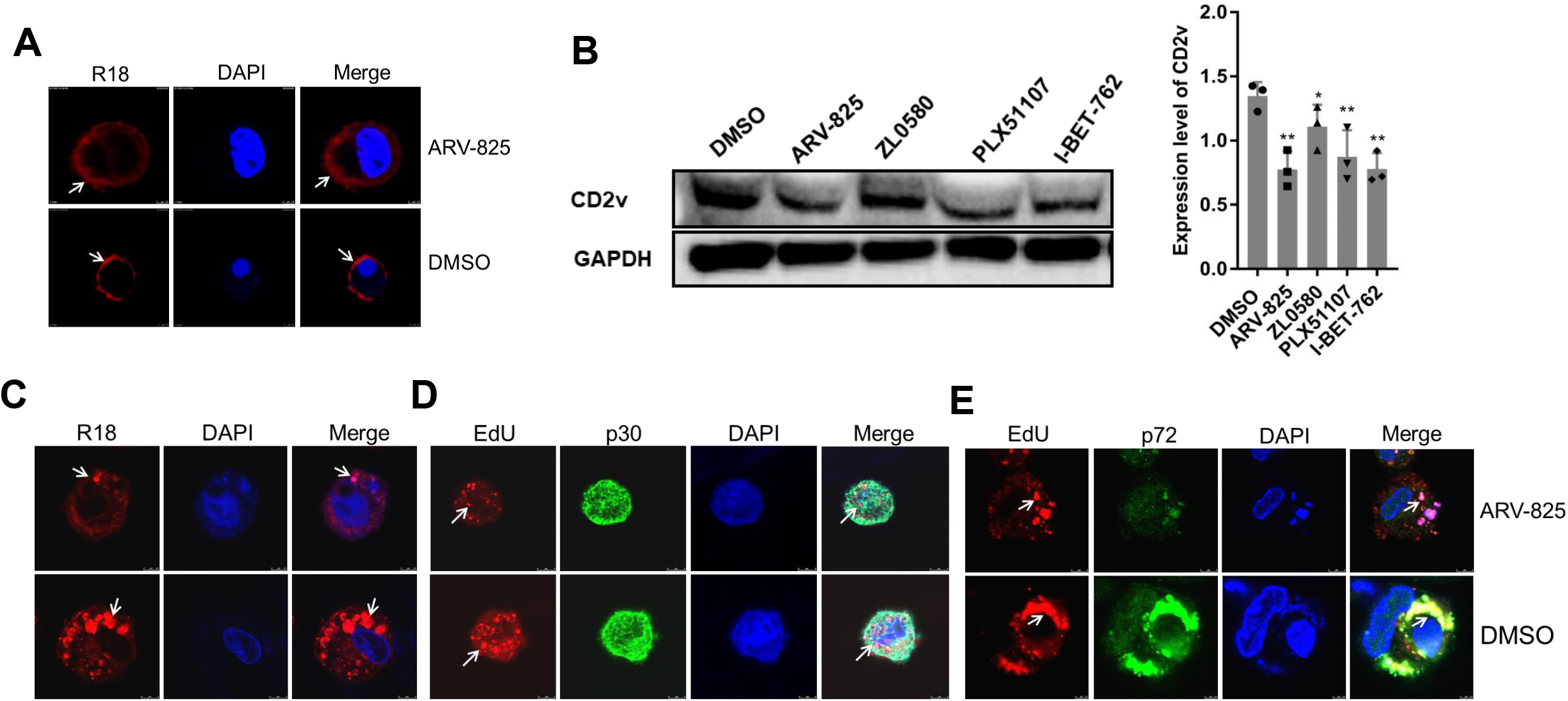
BRD4 inhibitors attenuate viral infection. (A) Viral attachment was assessed with fluorescence analysis in PAM cells treated with inhibitors and incubated with R18 labeled ASFV (MOI = 100). (B) Viral attachment was assessed with immunoblotting analysis against ASFV CD2v in PAM cells treated with inhibitors and incubated with ASFV (MOI = 10), GAPDH served as a loading control. (C) Viral internalization was assessed with fluorescence analysis in PAM cells treated with inhibitors and incubated with R18-labeled ASFV (MOI = 100). (D) Viral desencapsidation was assessed with fluorescence analysis in PAM cells treated with inhibitors and incubated with EdU-labeled ASFV (MOI = 100). (E) Viral factories were assessed with fluorescence analysis in PAM cells treated with inhibitors and incubated with EdU-labeled ASFV (MOI = 100). PAMs were seeded in 2-cm laser confocal dishes and pretreated with ARV-825 (0.5 μM) or DMSO at 37°C for 4h, and then removed the compounds. The PAMs were infected with ASFV and continued cultured at appropriate time points before collection.

### ZL0580, PLX51107, I-BET762 and ARV-825 suppress ASFV proteins synthesis

To further confirm the antiviral effects of ZL0580, PLX51107, I-BET762 and ARV-825, early and late expression of the important viral structural proteins p30 and p72, respectively, was analyzed by WB analysis (Figure 6A), Immunoblot analysis showed that in the presence of ZL0580, PLX51107, I-BET762 and ARV-825, the p30 and p72 protein levels in the PAMs were significantly reduced compared to those in the DMSO-treated cells, especially after treatment with ARV-825, and the expression levels of both proteins were decreased by more than 50%. Similar results were observed when expression of the ASFV p30 protein was evaluated by immunofluorescence analysis (Figure 6B). Clear fluorescent signals were detected, and the fluorescence density was higher in the DMSO-treated PAMs than in the inhibitor-treated PAMs. In contrast, the fluorescence intensity was significantly decreased in the 4 inhibitor treatment groups (Figure 6B). The percentage of cells showing early expression of the p30 protein was lower among cells treated with the BET inhibitors, as shown by flow cytometry analysis. ARV-825 treatment (1 μM) led to the sharply loss of p30 expression in ASFV-infected PAMs, with this effect followed by the effects of PLX51107, I-BET-762 and ZL0580 treatment. Compared with the DMSO-treated group, which was used as a control, the inhibitors had an at least 40% inhibitory effect (Figure 6C). These results indicated that BET/BRD4 inhibition affects the early and late protein synthesis.

**Figure 6:**
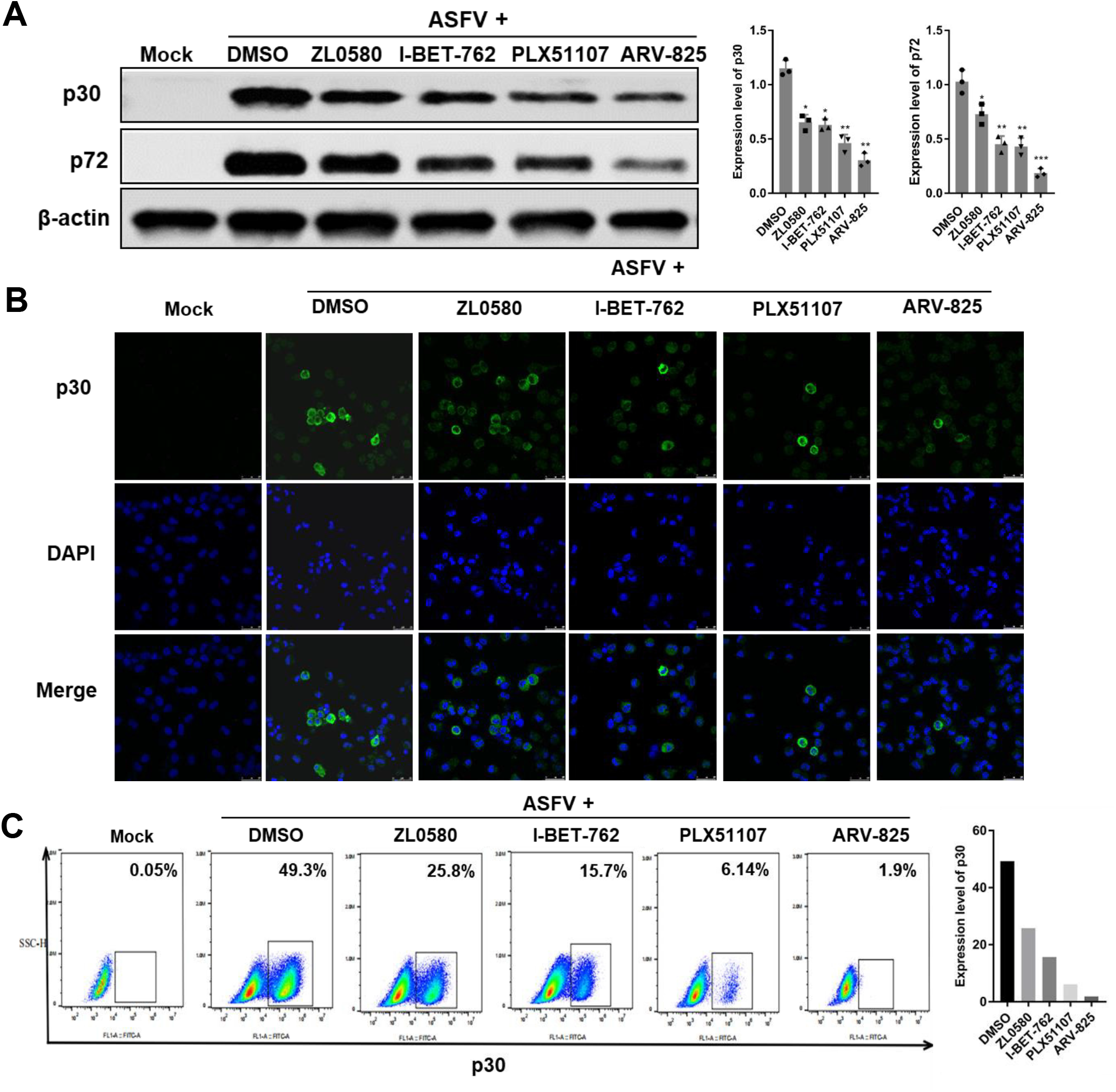
Evaluation of the inhibitory activity of ARV-825, I-BET-762, PLX51107 and ZL0580 against ASFV protein synthesis. (A) The expression of p30 and p72 in the presence of the 4 inhibitors ARV-825-1 μM, I-BET-762-10 μM, PLX51107-5 μM and ZL0580-10 μM was evaluated by WB analysis, (B) confocal microscopy and (C) flow cytometry. PAMs in 6-well plates were treated with inhibitors or DMSO 16 h prior to ASFV infection (MOI =1). The samples were collected at 24 h postinfection.

### BET/BRD4 inhibitors suppress the ASFV RNA polymerase expression levels

ASFV belongs to the nucleocytoplasmic large DNA virus (NCLDV) family, the members of which utilize quite complex RNA polymerases. Studies have shown that, unlike the 14 subunits encoded by eukaryotic RNA Pol II, ASFV encodes 9 subunits that are homologous to eukaryotic RNA Pol II subunits (20). BRD4 was proven to bind the positive transcription elongation factor (P-TEFb) to form a complex that is subsequently recruited to RNA Pol II of the host or virus, which then regulates the transcription of host or viral genes(17). Therefore, we further analyzed the transcription levels of the 9 subunits of ASFV RNA polymerase by real-time PCR. The results showed that ZL0580, PLX51107, I-BET-762 and ARV-825 significantly inhibited transcription of the ASFV RNA polymerase subunit genes compared to their transcription in the DMSO control group, and ARV-825 and ZL0580 treatment had a stronger inhibitory effect on gene transcription levels than treatment with the two other inhibitors (Figure 7A). We then selected ARV-825 and ZL0580 (BRD4-specific inhibitors) and evaluated their suppressive effects on the pC315R and pH359L proteins, which are homologs of TFIIB and RPB3 of the eukaryotic RNA Pol II (21), and the p30 protein. WB analysis revealed that ZL0580 and ARV-825-mediated inhibition of BET/BRD4 suppressed ASFV pC315R, pH359L and p30 protein expression (Figure 7B).

**Figure 7:**
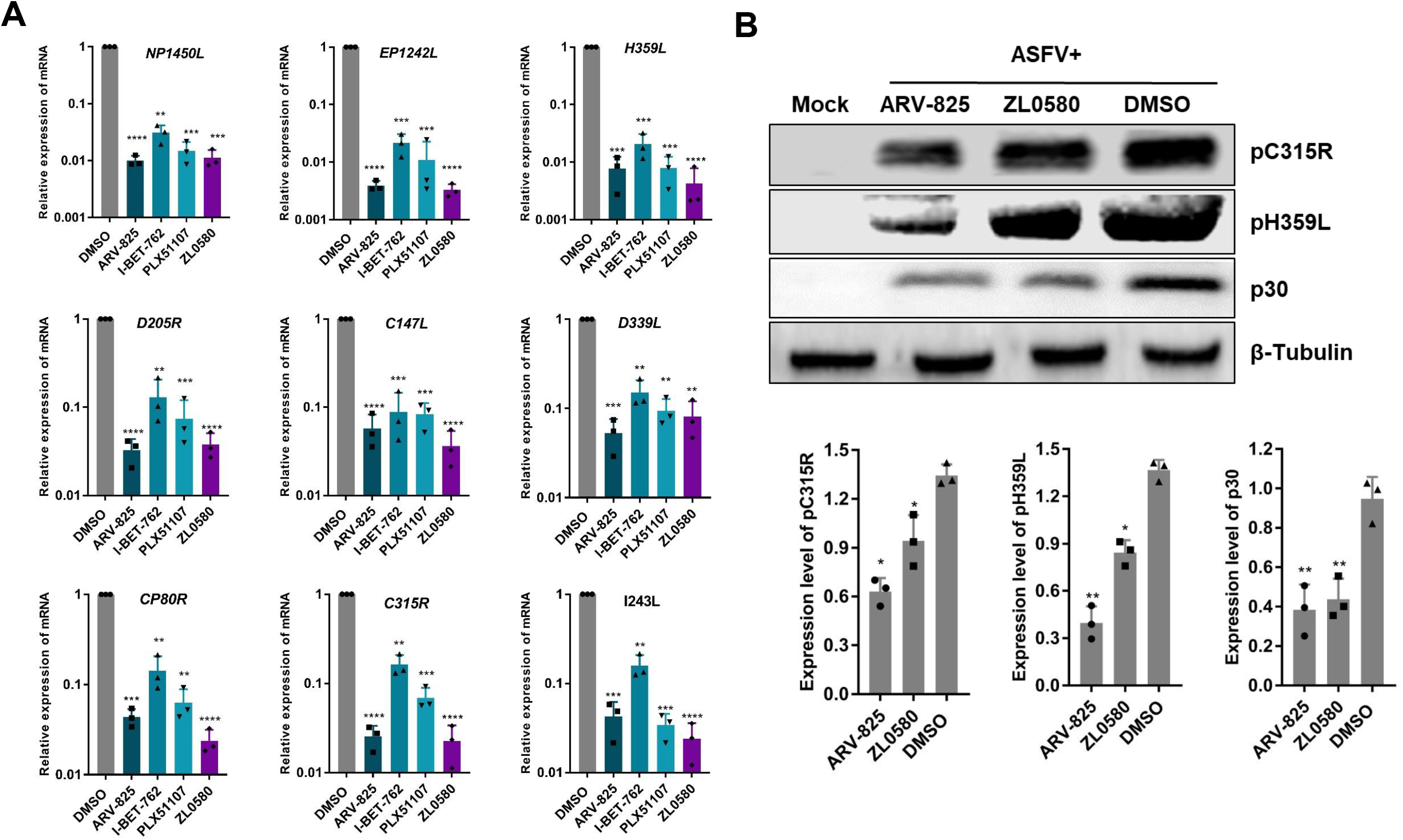
Effect of inhibitors on the putative subunits of ASFV RNA polymerase. (A) The expression of ASFV RNA polymerase subunits was significantly decreased at the RNA level and (B) protein level with inhibitor treatment. PAMs in 12-well plates were treated with ZL0580 (10 μM), I-BET-762 (10 μM), PLX51107 (5 μM), ARV-825 (1 μM) or DMSO for 16 h prior to ASFV infection (MOI =0.1), and the samples were collected at 24 h postinfection. The *NP1450L, EP1242L, H359L, D205R, C147L, D339L, CP80R, C315R* and *I243L* genes of ASFV were analyzed by qRT-PCR. The pC315R and pH359L proteins of ASFV were analyzed by WB analysis. Error bars show the SD of replicate qPCR experiments. All experiments were independently conducted at least 3 times. Statistical significance is denoted by \**P* < 0.05, \*\**P* < 0.01, \*\*\**P* < 0.001.

### Interaction with BRD4

We further analyzed 4 inhibitors’ biological effect on BRD4 (Figure 8). Docking analysis of BET-762, PLX51107, ARV-825 and ZL0580 binding to BD1 and BD2 domains of BRD4 was performed. BET-762 with BRD4 BD2 (PDB code: 5dfc). BET-762 interacts with Asn429 directly and forms three indirect hydrogen bonds with Tyr386, Leu381 and His433. PLX51107 with BRD4 BD1 (PDB code: 5wmg). PLX51107 interacts with Asn140 of BRD4 BD1 directly and forms indirect hydrogen bonds with Tyr97, Asp88 and Leu92. Besides, it has a salt bridge interaction with Lys91 via COOH and a π-π interaction with Trp81. OTX-015 (warhead of ARV-825) docked into BRD4 BD2. OTX-015 has a similar chemical structure with BET-762, thus their binding modes are resembling except the hydrogen bond with Asn429. ZL0580 docked into BRD4 BD1 and can’t be docked into the traditional KAc pocket only if the water molecules in the cavity were deleted. It interacts with Asn140 and Asp145 directly through hydrogen bonds.

**Figure 8:**
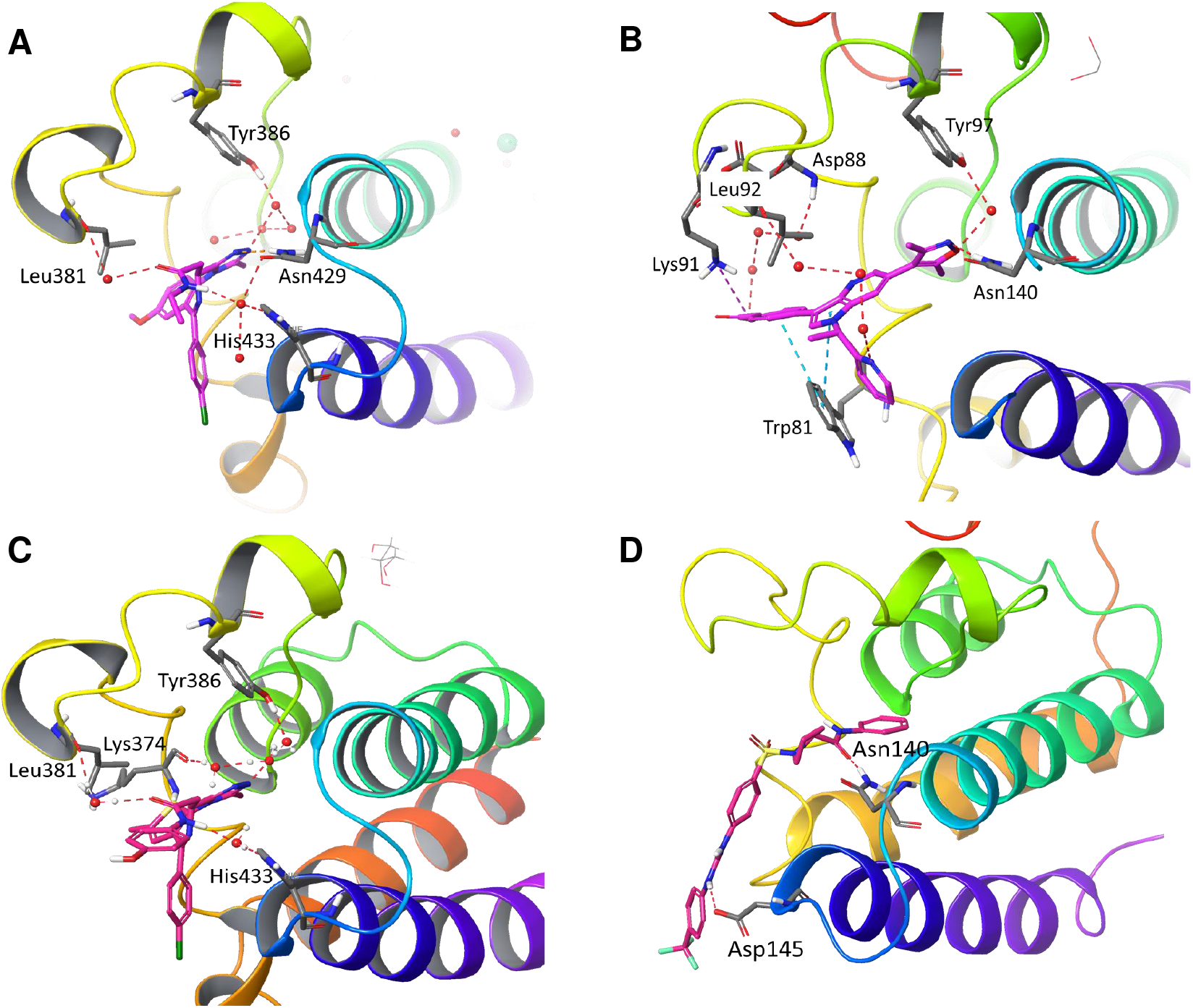
Interaction of 4 inhibitors with BRD4. Docking analysis of BET-762, PLX51107, ARV-825 and ZL0580 binding to BD1 and BD2 domains of BRD4 was performed with Schrödinger Small-Molecule Drug Discovery Suite.

## DISCUSSION

ASF, the most serious exotic pig disease, is listed as a class I animal disease in China. Since the first outbreak of ASF in Shenyang in August 2018 (22, 23), continuous infection has spread throughout the whole country, and ASF represents a serious threat for the global swine industry and the environment with grave economic consequences for stakeholders (24). The generation of vaccines can impede the global spread of ASF, in addition to the implementation of other measures, such as rapid diagnosis and control and eradication measures. However, commercialized vaccines for the prevention of ASFV infection remain lacking. In addition to vaccine development, the development of antiviral drugs is an important strategy to respond to ASF epidemics.

At present, the research and development of anti-ASFV drugs mainly focuses on two categories: (1) inhibitors that directly act on the proteins encoded by AFSV to affect its replication and (2) inhibitors that act on host protein factors required for viral replication to indirectly exert an anti-ASFV effect (25). Antiviral agents against ASFV currently include interferon (26), antibiotics (27), nucleoside analogues (28), plant-derived products (29) and other compounds that have been reported to inhibit ASFV replication (25). However, the safety of action of these antiviral drugs has not been studied in depth. Therefore, the need to identify new antiviral drugs for controlling ASFV is urgent.

Similar to other viruses, the signs of host infection with ASFV depend on the interaction between viruses and the host. BET family members include BRD2, BRD3, BRD4 and BRDT, which are widely involved in regulating the expression of genes related to transcription, DNA repair, immunity, metabolism and signal transduction; these proteins accomplish this by identifying acetylated histones or transcription factors via their two unique bromodomains and have become promising targets for tumor therapy and viral infection (10, 15). Small-molecule inhibitors of BET family proteins may provide a promising option for cancer treatment. To date, more than ten BET inhibitors have entered clinical trials and have mainly been used for the treatment of human diseases (11, 30). However, the effects of currently available BET/BRD4 inhibitors on ASFV infection are unknown.

During viral infection, host epigenetic factors can be involved in epigenetic modifications that affect the transcription and expression of viral genes and host genes (31, 32). Therefore, the elucidation of potential target genes of BET proteins may help reveal new functions of BET family members and provide new possibilities for clinical treatment and the combined application of BET inhibitors. The antiviral activity of BET inhibitors has been demonstrated against different viruses, including PRV (15), bovine papilloma virus (BPV) (19), human papilloma virus (HPV) (33), HIV (16), respiratory syncytial virus (RSV) (34) and Epstein-Barr virus (35). Previous studies have reported that BET inhibitors suppress the infectivity of these related viruses by decreasing macrophage and neutrophil infiltration into the airway, suppressing key inflammatory cytokines, preventing the expression of viral immediate-early proteins and/or effectively blocking BET/BRD4 phosphorylation-specific functions in transcription factor recruitment. Nevertheless, their antiviral effect on ASFV remains unknown.

In this study, we evaluated for the first time the antiviral effect of 12 representative BET/BRD4 inhibitors against ASFV infection *in vitro* (Figure 1). After screening for their cytotoxicity against PAMs by CCK-8 assay, 4 BET/BRD4 inhibitors were selected, and their roles in ASFV gene and protein expression were further studied. The cytotoxic effects of 12 BET/BRED4 inhibitors against PAMs were first evaluated by quantifying cell viability with a CCK-8 assay. Our results demonstrated that most of these BET inhibitors were less cytotoxic against PAMs at concentrations between 0.5 μM and 10 μM; therefore, we used doses of ≤10 μM (ARV-825: 1 μM; PLX51107: 5 μM; I-BET-762 and ZL0580: 10 μM) for further experiments. We determined the CC_50_ values for the 4 selected BET inhibitors to ensure their safety in PAMs (Figure 2B). In general, primary cells are more sensitive to compound cytotoxic effect than cell lines. However, in a previous study, obvious cytotoxicity was observed when the cells were treated with BET/BRD4 inhibitors (JQ-1, OTX-015 and I-BET 151) at 30 μM in both PK15 and HEK293 cells, while concentrations of 0-10 μM were minimally toxic in both cell lines (15), consistent with our results obtained with PAMs. This suggests that these inhibitors are harmless at concentrations below 10 μM in both primary cells and cell lines.

We performed time-of-addition studies to investigate whether the BET/BRD4 inhibitors have a primary antiviral effect on ASFV CN/SC/20109, a viral strain that replicates efficiently in primary PAMs (Figure 1). Early expression of the *CP204L* gene was significantly decreased when the inhibitors were applied prior to (pretreatment) or simultaneously with virus infection (*P* < 0.05), but the addition of inhibitors after ASFV infection (posttreatment) had no statistically significant effect on gene transcription levels. This suggests that the transcription of early viral genes is inhibited immediately by BET/BRD4 inhibitors when these genes begin to be largely transcribed. Interestingly, *B646L* gene expression was also obviously inhibited under cotreatment with all 12 inhibitors, but 2 inhibitors had no significantly effect on *B646L* gene transcription. Earlier addition of the inhibitors had a more notable inhibitory effect on ASFV, indicating that these 12 inhibitors act over the whole ASFV transcription process. Remarkably, two BET inhibitors (I-BET-762 and PLX51107) and two BRD4-specific inhibitors (ARV-825 and ZL0580) largely inhibited ASFV infection when applied in two ways (Figure 1A-B). Furthermore, the duration of action of I-BET-762, PLX51107, ARV-825 and ZL0580 was investigated in ASFV-infected cells for 4 h, 12 h, 24 h and 48 h (Figure 3). The cells were treated with individual inhibitors for 16 h prior to ASFV infection, and *B646L* gene transcription was then detected. The data indicated that ARV-825, I-BET-762 and PLX51107 suppressed *B646L* expression in ASFV-infected cells to a similar extent at 4 h and 12 h, but the effect became more significant when cells were infected at 48 h. For ZL0580, a stronger effect was observed in cells infected with ASFV for 4 h, and this effect even had a slight reduced when cells were infected with ASFV for 12 h, 24 h and 48 h. In addition, the cells were treated by BRD4 inhibitors prior to infected with ASFV at different time points dependent on the infected stages of its life cycle. The results indicated that the inhibitors could suppress ASFV infection. In general, the inhibitory effects were time-dependent. It is likely that BET/BRD4 inhibition induces cell cycle arrest or different biological activities; thus, the effects of these inhibitors on ASFV infection may vary by times being added to the culture.

Interestingly, some BET/BRD4 inhibitors do not affect PRV or PRRSV viral gene transcription (15), In contrast, other previous studies have demonstrated that BRD4 facilitates viral infection through the regulation of HSV-1 and HSV-2 viral gene transcription, and that through inhibiting BRD4, HSV-1 and HSV-2 viral infection and gene transcription, and protein synthesis were significantly suppressed in a dose-dependent manner (17). This suggests that modulation of similar or same target proteins (e.g., BET protein family) or pathways by different regulatory agents (different BET/BRD4 inhibitors) may induce distinct functional outcomes in different viral infections. Epigenetic modifications in the cell may be changed by altering activity of BET/BRD4 to further affect the transcription and expression of both viral and host genes. Understanding the regulatory mechanism of BET/BRD4 inhibitors and their roles in ASFV infection needs further studies. Collectively, our results suggest that BET inhibitors have therapeutic potential for control of ASFV infection.

I-BET-762 inhibits BET proteins by occupying the acetyl-lysine-binding pocket of BET proteins, inhibiting the binding of BET proteins to acetylated histones, and thereby prevents the formation of chromatin complexes responsible for the expression of key inflammatory genes in activated macrophages and primary human monocytes (36). PLX51107 is a novel, structurally distinct BET inhibitor. In a group of cultured cells, treatment with PLX51107 for a short period (4 hours) led to a sharp decrease in c-Myc levels but did not immediately cause an apoptotic response. After prolonged treatment time (continuous culture for 16 hours or longer), PLX51107 induced apoptosis. Proteolysis targeting chimeric (PROTAC) molecules are a novel family of compounds with the ability to bind their target proteins and recruit an ubiquitin ligase, which promotes degradation of the targeted protein (37). ARV-825 is a PROTAC compound and BRD4 protein degrader that can recruit BRD4 to the E3 ubiquitin ligase cereblon to induce rapid, effective and continuous degradation of the BRD4 protein, continuously lowering c-Myc levels (38, 39). Compared with other BRD4 inhibitors, ARV-825 treatment caused more significant changes in c-MYC levels and downstream cell proliferation and apoptosis induction (38). In our study, the dose-dependent inhibitory effects of ARV-825 were not as remarkable as those of ZL0580, PLX51107 or I-BET762. ARV-825 significantly inhibited ASFV *CP204L* and *B646L* mRNA and protein levels compared with those upon application of the other inhibitors at a lower concentration. To date, there have been no reports with respect to the effects of above three BET/BRD4 inhibitors on viral infection. ZL0580 is a BRD4-specific inhibitor that was designed by analyzing the crystal structures of available BRD4 modulators with the BRD4 BD1 domain (11). It displayed potent BRD4-binding activity with an IC_50_ = 163 nM against the BRD4 BD1 domain with 6.6-fold selectivity over the BRD4 BD2 domain. ZL0580 is a novel, BRD4-selective, small-molecule modulator that was reported to suppress both induced and basal HIV transcription and blocks viral reactivation events in human T cells and several latently infected myeloid cell lines. ZL0580 induces HIV suppression by inhibiting Tat-mediated transcription elongation and inducing a repressive chromatin structure at the HIV promoter (16, 40).

In our study, different assays (HAD, real-time PCR and WB analysis) showed that 4 inhibitors significantly inhibited ASFV infection in PAMs in a dose-dependent manner. A cumulative suppressive effect on ASFV infection was observed, suggesting that the BRD4 inhibitors specifically act on BRD4 to reduce ASFV infection (Figure 4-6). After characterizing these 4 inhibitors, we speculate that BET/BRD4 is helpful for ASFV infection, and the virus may take advantage of BET/BRD4 that is released from chromatin to the viral genome to promote viral replication and gene transcription. ASFV encodes approximately 20 genes that are involved in the transcription and modification of its mRNA (20). Our results indicated that the transcript levels of 9 related genes of ASFV were significantly decreased after treatment with the 4 individual BET/BRD4 inhibitors (Figure 7). ASFV carries a set of enzymes similar to eukaryotic RNA Pol II, and their homology with RNA Pol II is higher than that with other nuclear or cytoplasmic large DNA molecules (24). Interestingly, ZL0580, PLX51107, I-BET762 and ARV-825 inhibit the 9 subunits of ASFV RNA polymerase, which suggests that ZL0580, PLX51107, I-BET762 and ARV-825 exert their antiviral effects by altering ASFV transcription. It remains to be elucidated whether BET/BRD4 recruits P-TEFb to viral RNA polymerase and further regulates viral gene transcription or whether ASFV infection alters host chromatin status and utilizes host epigenetic modifications to facilitate viral replication. The regulatory mechanisms and the role of epigenetic BET/BRD4 in ASFV infection need to be further investigated.

While great progress has been made in understanding the interactions between ASFV and host, many unknowns still require further exploration. This study provides multiple lines of evidence to support that downregulation of early and late ASFV gene expression is associated with inhibition of BET/BRD4 activation and thus has a suppressive effect on ASFV infection. Extensive study of the role of BET/BRD4 in ASFV replication will be helpful to unravel the interactions between this virus and host cells and provide insights into development of new approaches for control of ASFV infection.

## MATERIALS AND METHODS

### Biosafety statement and facility

All experiments carried out with live ASFV were performed in a biosafety level-3 (BSL-3) laboratory at the Lanzhou Veterinary Research Institute (LVRI), Chinese Academy of Agriculture and Sciences (CAAS) and were accredited by the China National Accreditation Service for Conformity Assessment (CNAS) and approved by the Ministry of Agriculture and Rural Affairs. In the laboratory, to reduce any potential risk, all protocols were strictly followed, and all activities were monitored by the professional staff at LVRI and randomly inspected by local and central governmental authorities without advance notice.

### Cells culture and ASFV

Primary alveolar macrophages (PAMs) were isolated from 50-60-day-old specific pathogen-free (SPF) pigs and stored at the African Swine Fever Regional Laboratory (Lanzhou). The PAMs were cultured in RPMI 1640 medium (Thermo Scientific, USA) with L-glutamine and 25 mM HEPES (Gibco) supplemented with 10% fetal bovine serum (FBS, Gibco, Australia), 100 IU/ml penicillin and 100 μg/ml streptomycin (Gibco, Life Technologies) at 37 °C under 5% CO2. The ASFV strain used in this study (CN/SC/2019) was provided by the African Swine Fever Regional Laboratory (Lanzhou).

### Antibodies and reagents

For western blot (WB) analysis, anti-p30, -p72, anti-pC315R and anti-pH359L rabbit sera were raised against recombinant ASFV p30, p72 pC315R and pH359L proteins and deposited at the African Swine Fever Regional Laboratory (Lanzhou), Lanzhou Veterinary Research Institute (LVRI) of the Chinese Academy of Agricultural Sciences. Anti-CD2v mouse sera were kindly provided by Prof. Liguo Zhang from Institute of Biophysics, Chinese Academy of Sciences. Anti-β-Actin (13E5) rabbit monoclonal antibody (Cat. no. 4970) and anti-GAPDH (14C10) rabbit monoclonal antibody (Cat. no. 2118) were purchased from Cell Signaling Technology. Anti-β-tubulin rabbit polyclonal antibody (Cat. no. 10094-1-AP) and HRP-conjugated AffiniPure goat anti-rabbit IgG (H+L) (Cat. no. SA00001-2) were purchased from ProteinTech Group. FITC-conjugated goat anti-rabbit IgG secondary antibody (Cat. no. F0382) was purchased from Sigma-Aldrich. A Cell Counting Kit-8 (CCK-8) (Cat. no. K1018-30) was purchased from APExBIO (MA, USA). TRIzol™ reagent (Cat. no.15596018), 4’, 6-diamidino-2’-phenylindole (DAPI) (Cat. no. 62248), fluorescent dyes R18 (Cat. no. O246) and RIPA lysis and extraction buffers (Cat. no 89901) were purchased from Thermo Fisher Scientific. 5-ethynyl-2’-deox-yuridine (EdU) (Cat. no C00031) was purchased from Guangzhou RIBOBIO. Co., Ltd.

### BET/BRD4 chemical inhibitors

Apabetalone (RVX-208) (BET inhibitor, S7295), ARV-825 (BRD4 specific inhibitor, S8297), AZD-5153 6-hydroxy-2-naphthoic acid (BET/BRD4 inhibitor, S8344), CPI-203 (BET inhibitor, S7304), Molibresib (I-BET-762) (BET inhibitor, S7189), (+)-JQ1 (BET inhibitor, S7110), INCB054329 (BET inhibitor, S8753), MS436 (BET inhibitor, S7305), Birabresib (OTX015) (BET inhibitor, S7360), PLX51107 (a new BET inhibitor, S8739) and PFI-1 (PF-6405761) (BRD2/BRD4 inhibitor, S1216) were purchased from Selleck.cn, ZL0580 (BRD4 specific inhibitor) was prepared as previously described (16). The structures and functions of these inhibitors are shown in Figure 2A and S1 (https://www.selleck.cn/) and Table 1 (16, 38, 41-50).

**Table 1.** BET/BRD4 chemical inhibitors used in this study.

| Inhibitors | Functions | References |
| --- | --- | --- |
| Apabetalone (RVX-208) | The effective BET bromodomain inhibitor, act on BD2. | (41) |
| ARV-825 | Recruit BRD4 to E3 ubiquitin ligase cereblon to induce rapid, effective and continuous degradation of BRD4. | (38) |
| AZD5153 | BET/BRD4 bromodomain BD2 inhibitor, inhibit the target gene expression of nuclear receptor binding SET domain protein 3 (NSD3). | (42) |
| CPI-203 | Potent BET bromodomain inhibitor. | (43) |
| Molibresib (I-BET-762) | A highly selective inhibitor of BET family. | (44) |
| INCB-054329 | BET family bromodomain inhibitor. | (45) |
| (+)-JQ1 | BET bromodomain inhibitor, (+)-JQ1 inhibits cell proliferation by inducing autophagy; (+)-JQ1 can inhibit the target gene expression of nuclear receptor binding SET domain protein 3 (NSD3). | (46) |
| MS436 | BET bromodomain inhibitor. | (47) |
| Birabresib (OTX015) | Specifically binds to BRD2/3/4, inhibit the target gene expression of nuclear receptor binding SET domain protein 3 (NSD3). | (48) |
| PLX51107 | A novel BET inhibitor. Among non-BET proteins, PLX51107 only has a significant interaction with the bromine region of CBP and EP300 (p300). | (49) |
| PFI-1 (PF-6405761) | A highly selective BET inhibitor that acts on BRD4 and BRD2. | (50) |
| ZL0580 | ZL0580 is selectively bound to the BRD4 BD1 domain that induced epigenetic suppression of HIV via BRD4. | (16) |

### Cytotoxicity assay

The cytotoxicity of 12 representative inhibitors in PAMs was evaluated by using a CCK-8 kit. Briefly, PAMs (2 × 10^5^ cells per well) in 96-well cell culture plates were treated with the inhibitors at increasing concentrations (from 0.5 μM to 240 μM). The experiments included three replicates, and a blank and DMSO control were included. The treated cells were incubated for 24 h at 37 °C in 5% CO_2_, and after incubation, 10 μL of CCK-8 reagent was added to each well and incubated at 37 °C for 1-4 h. The absorbance at 450 nm was measured using a microplate reader. The viability of the PAMs was calculated according to the following formula: cell viability (%) = [(OD inhibitor - OD blank) / (OD control- OD blank)] × 100.

### Virus HAD_50_ assay

PAMs were seeded in 96-well plates and cultured overnight at 37 °C under 5% CO2. The cells were then pretreated with DMSO, PLX51107, ARV-825, ZL0580 and I-BET-762 for 16 h, after which 10-fold serial dilutions (10^0^-10^−12^) of virus were inoculated into each well (with eight replicates for each dilution), with pig erythrocytes (1:1000) added to each well at the same time. The ASFV was quantified by the formation of characteristic rosettes formed through hemadsorption (HAD) of erythrocytes around the infected cells. HAD activity was observed for 7 consecutive days after inoculation, and the 50% HAD dose (HAD_50_) was calculated by using the Reed and Muench method (51).

### Time-of-addition assay

PAMs in 12-well plates (2 × 10^6^ cells/well) were seeded for ASFV infection. In the pretreatment assay, PAMs were treated with 12 individual BET/BRD4 inhibitors for 16 h before infection with ASFV CN/SC/2019 (MOI = 0.1). In the cotreatment assay, PAMs were exposed to 12 individual BET/BRD4 inhibitors at the same time that the ASFV was added to the plates. The plates were then incubated at 37 °C under 5% CO_2_ for 24 h. In the posttreatment assay, cells were infected with ASFV, and the inhibitors were then added 4 h after infection. The plates were then incubated at 37 °C under 5% CO_2_ for 16 h. DMSO-treated cells when then infected with ASFV for different assays in individual wells. The viruses were collected, titrated by HAD assay, and quantified by real-time PCR and WB analysis.

### Quantification of cell-associated ASFV mRNA

To quantify ASFV mRNA in ASFV-infected PAMs, total RNA was extracted from different PAM samples using a standard protocol with TRIzol™ reagent (Life Technologies), followed by chloroform extraction and precipitation with isopropyl alcohol and ethanol. cDNA was synthesized from the RNA using the PrimeScriptTM RT Reagent Kit with gDNA Eraser (Takara Bio Inc, Shiga, Japan) according to the manufacturer’s instructions. Gene expression in the cDNA samples was measured by one-step qRT-PCR using a One Step PrimeScript RT-PCR Kit (Perfect Real Time) according to the manufacturer’s specifications (Takara, Dalian, China). Quantitative real-time PCR was performed on the CFX Connect Real-Time PCR Detection System (Bio-Rad, USA). The sequences of primers and probes specific for the *B646L* gene were obtained according to the OIE-recommended sequence described in King et al (52). Primer and probe sequences specific for other genes were designed in this study (Table 2). All samples were run and analyzed in duplicate. The RNA expression of each target gene in the PAMs was normalized to GAPDH expression and then calculated using the 2^- ΔΔ^CT method.

**Table 2.**
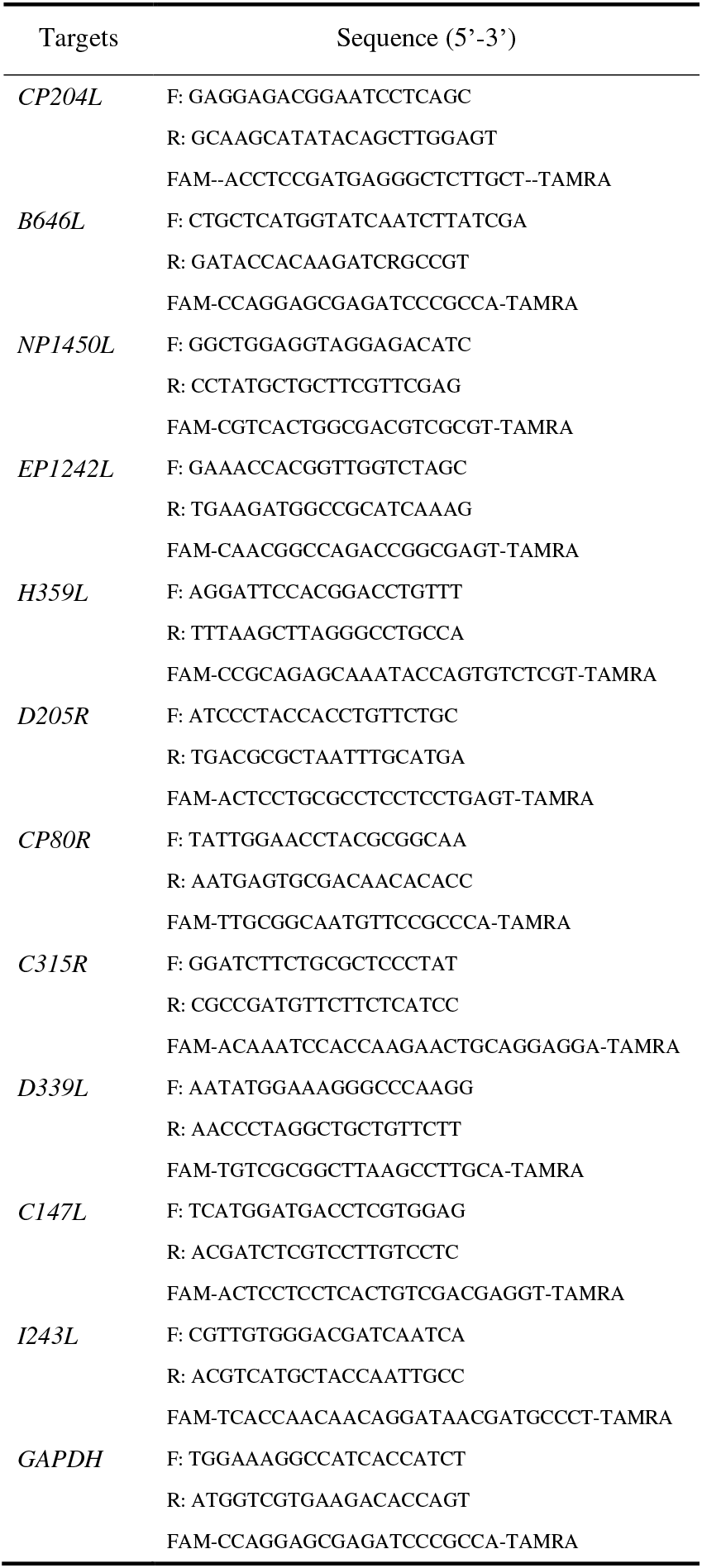
List of primers and probes used in this study.

### Western blotting (WB) analysis

PAMs were seeded in 6-well plates overnight and treated with ZL0580, I-BET-76, PLX51107, ARV-825 or DMSO 16 h prior to inoculation, followed by infection with the ASFV CN/SC/2019 strain (MOI=1) for 48 h. The cells were harvested, washed and then lysed in RIPA lysis and extraction buffers supplemented with a protease inhibitor cocktail and 1 mM PMSF by rotation at room temperature (RT) for 1 h. The total protein concentration was measured using a Microplate BCA Protein Assay Kit (Pierce™, Thermo Fisher Scientific). Proteins were separated on an SDS-PAGE gel and then transferred to a nitrocellulose (NC) membrane (Merck Millipore, ISEQ00010), which was incubated with individual protein-specific primary antibodies at 4 °C overnight on a shaker. The membrane was then incubated with horseradish peroxidase (HRP)-linked secondary antibodies for 1 h at RT. The reaction was detected with Immobilon™ western HRP substrate (B1911-100ML, Sigma). The corresponding grayscale value for each expressed protein band was analyzed using ImageJ software.

### Confocal microscopy

PAMs were seeded in 2-cm laser confocal dishes and pretreated with ZL0580 (10 μM), I-BET-762 (2 μM), PLX51107 (5 μM), ARV-825 (0.5 μM) or DMSO at 37°C for 4h, and then removed the compounds. The treated PAMs were infected with ASFV or fluorescent dye R18 labeled ASFV at an MOI of 100 at 4 °C for 1 hour (binding), or cultured the cells at 37 °C for 1.5h (internalization), or ASFV-infected cells were exposed to EdU for 3h (desencapsidation) or 24h (viral factories) before collection. The cells were fixed in a buffer containing 4% paraformaldehyde and 10 mM piperazine-N, N-bis (2-ethanesulfonic acid) in PBS at pH 6.4 for 10 min. After one wash and incubation with primary antibodies diluted in blocking buffer without Triton X-100 at 4 °C overnight. Then, the cells were stained with FITC-conjugated goat anti-rabbit IgG secondary antibody for 1 h at RT. Cells were acquired using confocal microscopy (Leica, TCS SP8).

### Flow cytometry assay

Cells were pre-treated with compounds for 16 h, and then infected with ASFV (MOI = 1) and simultaneously treated with compounds for 24 h. Cells were collected by centrifugation and suspended in phosphate-buffered saline (PBS) and stained for DNA (with DAPI), and anti-ASFV rabbit antibody p30. Cytometry acquisition was performed on a BD Accuri® C6 Plus instrument, and the data were analyzed using the program FlowJo v10.6.2.

### Binding modes of PLX51107, I-BET762, ZL0580 and ARV-825 with BRD4

The docking study was performed with Schrödinger Small-Molecule Drug Discovery Suite. The crystal structure of BRD4 BD1 (PDB code: 5wmg) and BRD4 BD2 (PDB code: 5dfc) were downloaded from RCSB PDB Bank and prepared with Protein Prepared Wizard. During this step, hydrogens were added, crystal waters were removed while water molecules around the KAc pocket were kept (all the water molecules were removed in the case of ZL0580), and partial charges were assigned using the OPLS-2005 force field. The 3D structures of ZL0580 and ARV-825 were created with Schrödinger Maestro, and the initial lowest energy conformations were calculated with LigPrep. For all dockings, the grid center was chosen on the centroid of included ligand of PDB structure KAc site and a 20 × 20 × 20 Å grid box size was used. All dockings were employed with Glide using the SP protocol. Docking poses were incorporated into Schrödinger Maestro for a visualization of ligand-receptor interactions.

### Statistical analysis

Statistical analyses of all data were performed using Prism 8.0 (GraphPad Software, Inc.). Statistical comparisons between groups were performed using paired or nonpaired t tests. Two-tailed p values were determined, and a p value < 0.05 was considered to indicate statistical significance (* *P* < 0.05; ** *P* < 0.01, *** *P* < 0.001 and **** *P* < 0.0001). The quantitative data in all Figures are expressed as the mean ± SD (indicated by the error bars). The 50% cell cytotoxicity (CC_50_) was calculated by a linear regression analysis of dose-response curves generated from the obtained data. The 95% confidence intervals (95% CIs) for CC_50_ values were calculated using IBM SPSS Statistics v19.0.

## ACKOWLEDGEMENTS

This study was financially supported by the National Natural Science Foundation of China (Grant Nos.32072830); Gansu Provincial Major project for science and technology development (Grant Nos. 20ZD7NA006); State Key Laboratory of Veterinary Etiological Biology, Lanzhou Veterinary Research Institute, Chinese Academy of Agricultural Sciences (Grant Nos. SKLVEB2020CGPY02); Basic scientific research business expenses budget incremental project, Chinese Academy of Agricultural Sciences, Lanzhou veterinary research institute (Grant Nos. 1610312021002).

QLN, GQG, ZJL, JXL and HY conceived and designed the study, YRZ, SXY and QLN participated in the whole experiments, QLN wrote the manuscript. JJY, ZHZ, SXG and JF isolated the PAM cells. ZQL and JZ synthesized the BRD4 inhibitor ZL0580 and revised the manuscript. HTH analyzed the data and revised the manuscript.

We would like to thank Dr. Yongxiang Fang from Lanzhou Veterinary Research Institute (LVRI), Chinese Academy of Agriculture and Sciences (CAAS) for help with the flow cytometry assay; AJE for English language editing. All authors reviewed the manuscript and declared that they have no competing interests.

